# Pathway and gates for ATG2A-mediated lipid transport in autophagy

**DOI:** 10.1101/2025.11.16.688672

**Authors:** Ainara Claveras Cabezudo, Elisabeth Holzer, Sascha Martens, Gerhard Hummer

**Affiliations:** Max Planck Institute of Biophysics, Department of Theoretical Biophysics, Frankfurt am Main; IMPRS on Cellular Biophysics, Frankfurt, Germany; Institute of Biophysics, Goethe University Frankfurt, Frankfurt, Germany; Max Perutz Labs, Vienna Biocenter Campus (VBC), Vienna, Austria; University of Vienna, Max Perutz Labs, Vienna, Austria; Vienna Biocenter PhD Program, a Doctoral School of the University of Vienna and the Medical University of Vienna, Austria; Aligning Science Across Parkinson’s (ASAP) Collaborative Research Network, Chevy Chase, USA

## Abstract

Autophagy is a complex process in which eukaryotic cells degrade cytosolic components by delivering them to lysosomes via double-membrane autophagosomes. The lipid transfer protein ATG2A plays a crucial role in autophagosome formation by tethering the phagophore membrane to the ER and delivering a significant fraction of the required lipids. The mechanism by which ATG2A shuttles lipids from one membrane to the other, however, remains elusive. Here, we combine structural predictions, molecular dynamics simulations and *in vitro* lipid transfer assays to gain mechanistic insights into ATG2A-mediated lipid transport. Using this integrative approach, we characterize the contact sites of the protein with donor and acceptor membranes. Our simulations capture multiple events of lipid uptake and delivery from and to the bound membrane. Conformational rearrangements of N-terminal amphipathic helices emerge as a critical factor for facile lipid transport. With this insight, we designed an ATG2A mutant that transfers lipids three times faster than the wild type *in vitro*. In complex with ATG9A, ATG2A forms a bridge between two parallel membranes at ⍰12 nm separation. Overall, our findings suggest that ATG2A is a lipid transporter gated at the N-terminus by blocking helices that, upon release, act as additional membrane tethers.

## 1 Introduction

Autophagy is a complex molecular process in which eukaryotic cells degrade cellular components and recycle the building blocks. During this process, a double-membrane organelle named autophagosome engulfs a part of the cytosol and delivers its content to the lysosome (or the vacuole in yeast) for degradation.^16,18,23,24^ This mechanism plays an essential role under stress conditions and is crucial to maintain cell homeostasis.^9^ Defects in autophagy have been linked to several neuropathologies, including Alzheimer’s, Parkinson’s and Huntington’s disease.^8,25,37^ Moreover, autophagy has been shown to play important roles in tumor initiation and progression,^9^ which makes it a very promising therapeutical target.

Atg2, a core autophagy protein conserved from yeast to human, plays a crucial role in phagophore expansion. Over the past years, several studies have shown that Atg2 transfers lipids between vesicles *in vitro* and is crucial for phagophore elongation *in vivo*.^20,28^ Taken together, these data suggest that Atg2 mediates phagophore expansion by tethering the growing phagophore to the ER and providing a substantial number of the required lipids.

Despite the crucial role of Atg2-mediated lipid transport in autophagy, important aspects of this mechanism remain elusive. These include the driving force and specificity of the transport, as well as the orientation and contact sites of Atg2 with respect to the donor and acceptor membranes. In addition, lipid transfer experiments have shown that the transport exhibits a slow and a fast phase,^20,28,36^ whose nature is not understood.

Atg2 has multiple interaction partners. One of them is the lipid scramblase Atg9, which is the only transmembrane protein of the autophagy machinery.^19,21,27^ Atg9 has been reported to interact with the C-terminal region of Atg2 at the phagophore site and equilibrate the two leaflets of the acceptor membrane.^12^ The lipid scramblases TMEM41B and VMP1, localized in the ER, interact with the N-terminal region of ATG2A,^11^ the human ortholog of Atg2 in yeast. WIPI4, as one of the four human orthologues of Atg18 in yeast, also interacts with the C-terminus of ATG2A and has been proposed to have a recruiting role to the phagophore.^7^ The extent to which the Atg2-mediated lipid transport depends on its interaction partners, however, remains unclear.

Over the last few years, several efforts to resolve the structure of Atg2 alone and with its interaction partners have been conducted. In a seminal study, Osawa and colleagues obtained an X-ray structure of the N-terminus of yeast Atg2 (*S. pombe*), shedding light on the mechanism of lipid uptake.^28^ Very recently, Wang et al.^38^ have resolved a cryo-EM structure of the human homologue ATG2A in complex with WIPI4 and with ATG9A. van Vliet et al.had previously proposed an alternative model of the ATG2A-ATG9A complex.^34^ Both models support a direct handover of lipids from ATG2A to ATG9A, yet differ on the exact delivery site. Despite these advances, the dynamic nature of lipid transport presents a major challenge. For instance, the N- and C-terminal regions of ATG2A are not resolved in the cryo-EM structure obtained by Wang et al.^38^ due to their high flexibility. These regions, however, are crucial for lipid uptake and delivery. Moreover, the structures were determined in the absence of lipid bilayers, which may influence the structure and dynamics of proteins.

In this study, we combine structural predictions, coarse-grained (CG) molecular dynamics (MD) simulations, and *in vitro* lipid transfer assays to gain mechanistic insights into lipid transport mediated by ATG2A. Unbiased simulations reveal interactions between the N- and C-terminal sites of ATG2A and the membrane. We observe lipid uptake and delivery from and to lipid aggregates, supporting the idea that ATG2A functions as a self-sufficient lipid transporter. Our findings, supported by experimental evidence, suggest that rearrangements in the N-terminal region are critical for lipid uptake from the ER, pointing towards a potential regulatory mechanism. Finally, we propose a structural model of the ATG2A-ATG9A complex which does not involve direct lipid delivery from ATG2A to ATG9A.

## 2 Results

### 2.1 ATG2A binds lipids at both termini in an orientation compatible with lipid transport

We used AlphaFold 3^2^ to build a structural model of the full-length human ATG2A protein that includes the C- and N-terminal regions, which have so far remained elusive to structural studies due to their high flexibility.^38^ The predicted structure includes the previously characterized N-terminal chorein N-like domain, the rod-shaped hydrophobic cavity, and a C-terminal region that contains amphipathic helices. Notably, the hydrophobic groove is accessible through the C-terminus, but not through the N-terminus (Figure 1a).

**Figure 1.**
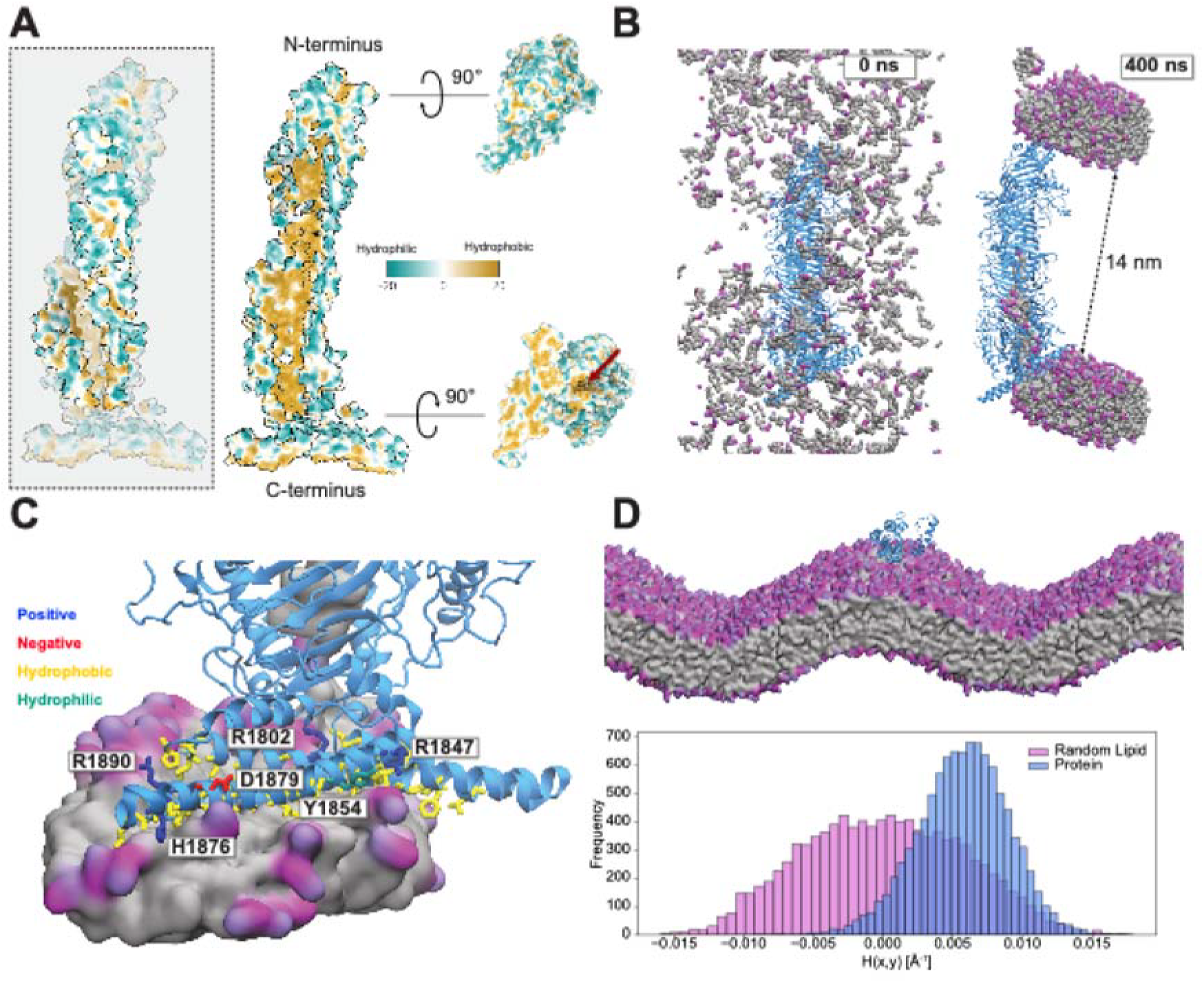
ATG2A binds two parallel lipid bilayers at its termini. (A) The hydrophilic outside (blue) of ATG2A (left) contrast with a long hydrophobic cavity (gold) visible in a cut along the principal axis (right). The cavity is accessible from the C-terminus (bottom), but not from the N-terminus (top). (B) First (left) and last frame (right) of an MD simulation with initially dispersed POPC lipids that then self-assemble and interact with both ends of ATG2A, accessing the cavity through the C-terminal end. (C) Interactions between the C-terminus (cartoon) and a lipid cluster (surface render). Hydrophobic residues interact with the lipid tails (gray), while charged and hydrophilic residues engage in interactions with the head groups (magenta). (D) The C-terminus of ATG2A has a preference for positively curved bilayers. Snapshot of a simulation of the C-terminus of ATG2A on a buckled bilayer (top) and mean bilayer curvature at the center of mass of the protein (bottom) compared to the one of a random lipid. For each frame along the trajectory, a lipid was randomly selected and the local curvature at the position of its phosphate head group was calculated. The histograms show the curvature values of 10 independent replicas, each of 1 µs of duration. All snapshots correspond to CG simulations after backmapping the protein.

To characterize the protein-lipid contact sites in an unbiased manner, we ran coarse grained (CG) simulations of ATG2A with 1-palmitoyl-2-oleoyl-sn-glycero3-phosphocholine (POPC) lipids that were initially dispersed in solution (Supplementary Table 1, Methods). After a few nanoseconds, lipids started to self-assemble to form micelles and then bicelle-like structures, some of which localized to the N- and C-termini of ATG2A (Supplementary Movie 1). In most simulations, the two bicelles at the N- and C-termini were nearly parallel to each other, with a midplane-midplane distance of ⍰ 18 nm and an edge-edge distance of ⍰ 14 nm (Figure 1b). Similar distances and membrane orientations have been reported *in vitro* and *in vivo*.^6,7^

ATG2A has been shown to tether vesicles in different ways: while the lipid shuttle preferentially engages with vesicles in a rather perpendicular orientation, a small subpopulation of ATG2A binding along the membrane tangent in a flat orientation has been observed using cryo-ET.^38^ In agreement with these findings, we observe transient interactions of micelles with some of the disordered regions at the surface of the hydrophobic groove, sometimes with two micelles binding simultaneously at opposing sides (Supplementary Figure 1).

In our MD simulations, interactions between lipid bicelles and the amphipathic helices at the C-terminal region of ATG2A were remarkably stable. These interactions involve hydrophobic residues, which engage with the lipid tails, and hydrophilic and charged residues, which interact with the charged phosphate and choline head groups (Figure 1c). Although interactions between lipids and the N-terminus are also relatively frequent, these are more short-lived in the MD simulations (Supplementary Movie 1).

We noticed that the small bilayer-like lipid assemblies forming at the C-terminus of ATG2A in our simulations frequently exhibited high curvature. To elucidate whether the amphipathic helices at this end have a membrane-curvature preference, we ran simulations of this region of the protein (residues 1-239) on a buckled lipid bilayer. These simulations show a strong preference of the C-terminus of ATG2A for positively curved regions of the membrane (Figure 1d, Supplementary Movie 2). This observation is in agreement with previously reported experimental evidence that Atg2 preferentially binds to highly curved membranes with lipid packing defects.^7,12,15,28^

### 2.2 ATG2A mediates bidirectional transport of lipids

In multiple MD simulations, the C-terminal site of the hydrophobic cavity of ATG2A is filled up with one or several lipids. Some of these lipids enter into the cavity through a lateral opening which has been observed in the cryo-EM structure of ATG2A^38^ (Supplementary Movie 1) or are randomly placed at the cavity during system set-up. In addition, we also observed lipid uptake from micelles through the C-terminal end of the cavity, indicating that our ATG2A structural model is transport competent. Interestingly, we only observed lipid uptake when the cavity contained at least one other lipid. The process of lipid entry resembles the flip-flop mechanism of lipids in membranes,^3,30^ with one tail accessing the cavity first and “dragging” the rest of the lipid inside (Figure 2a, Supplementary Movie 3). Lipid release from the C-terminus into micelle-like lipid aggregates was also observed during the simulations, indicating that lipid transport at the C-terminus is bidirectional (Figure 2b, Supplementary Movie 4).

**Figure 2.**
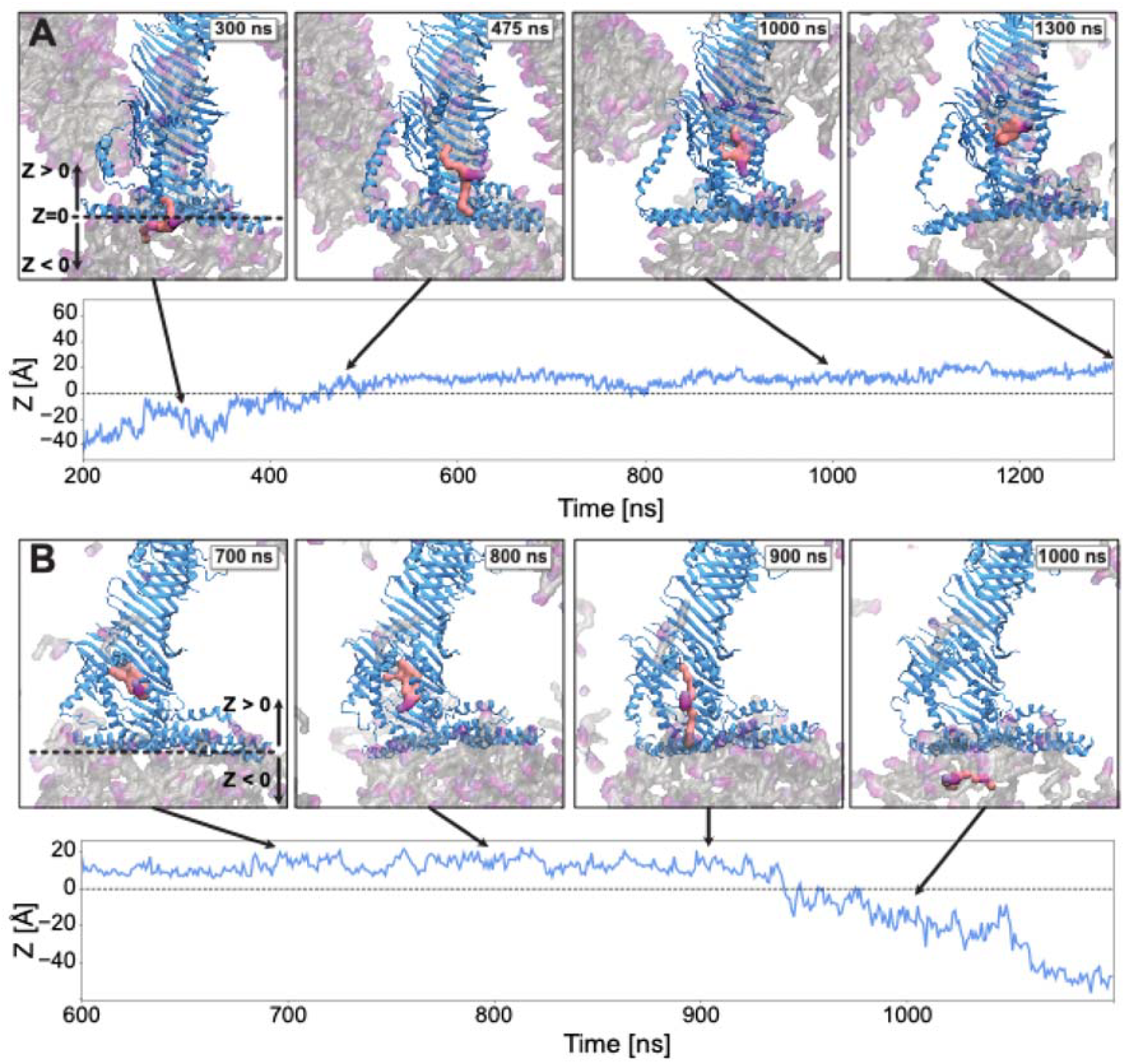
Bidirectional transport of lipids at the C-terminus of ATG2A. (A) Lipid (pink) entry from a micelle (grey tails with magenta headgroups) into the C-terminal cavity. (Bottom) Position of the phosphate head group along the *Z*- axis as function of time, with the entry point of the hydrophobic cavity set at *Z* = 0. (B) Lipid exit from the C-terminal cavity of ATG2A into a micelle. All snapshots correspond to CG simulations after backmapping the protein. The vertical position *Z* of the lipid relative to the cavity entry point is shown as function of time. Disordered loops in the structure are omitted for clarity.

Although the cavity is not accessible for lipid uptake through the N-terminus, it could in principle be filled up with lipids entering through the C-terminus, forming a continuous lipid channel. In our simulations, we observed, however, a maximum of 10 lipids in the cavity. Lipids consistently filled up the cavity up to a specific point located ⍰ 6.5 nm away from the C-terminal entry, pointing towards a barrier for lipid passage in this region (Supplementary Figure 2a).

In the cryo-EM structure of ATG2A resolved by Wang et al.,^38^ residues L603 and F678 interact, locally tightening the hydrophobic cavity at the middle passage region. We thus monitored these residues during our simulations, and observed that L603 and F678 are located in a comparably narrow region that is typically not occupied by lipids (Supplementary Figure 2a). Surprisingly, however, in one simulation a lipid, which in the initial random setup had been placed at the cavity, jumped to this site and remained there for hundreds of nanoseconds (Supplementary Figure 2b, Supplementary Movie 5). This suggests that the middle passage region can stably accommodate lipids and may play a role in regulating lipid passage through the cavity.

### 2.3 A rearrangement of N-terminal amphipathic helices is necessary for lipid transport

In the AlphaFold 3 model of ATG2A, the cavity is not accessible for lipids through the N-terminus (Figure 1a), with three *α*-helices blocking the cavity. One of these helices is at the very N-terminus (H1, residues 13-30), and the other two are downstream in the sequence and connected by a short linker (H2, residues 125-142 and H3, 151-168). These helices are predicted to be amphipathic (Figure 3a). It has been proposed that a displacement of the helix H4 in yeast ATG2A is required for phospholipid uptake.^28^ We thus hypothesized that rearrangements of these three helices in the human ATG2A might take place as well.

**Figure 3.**
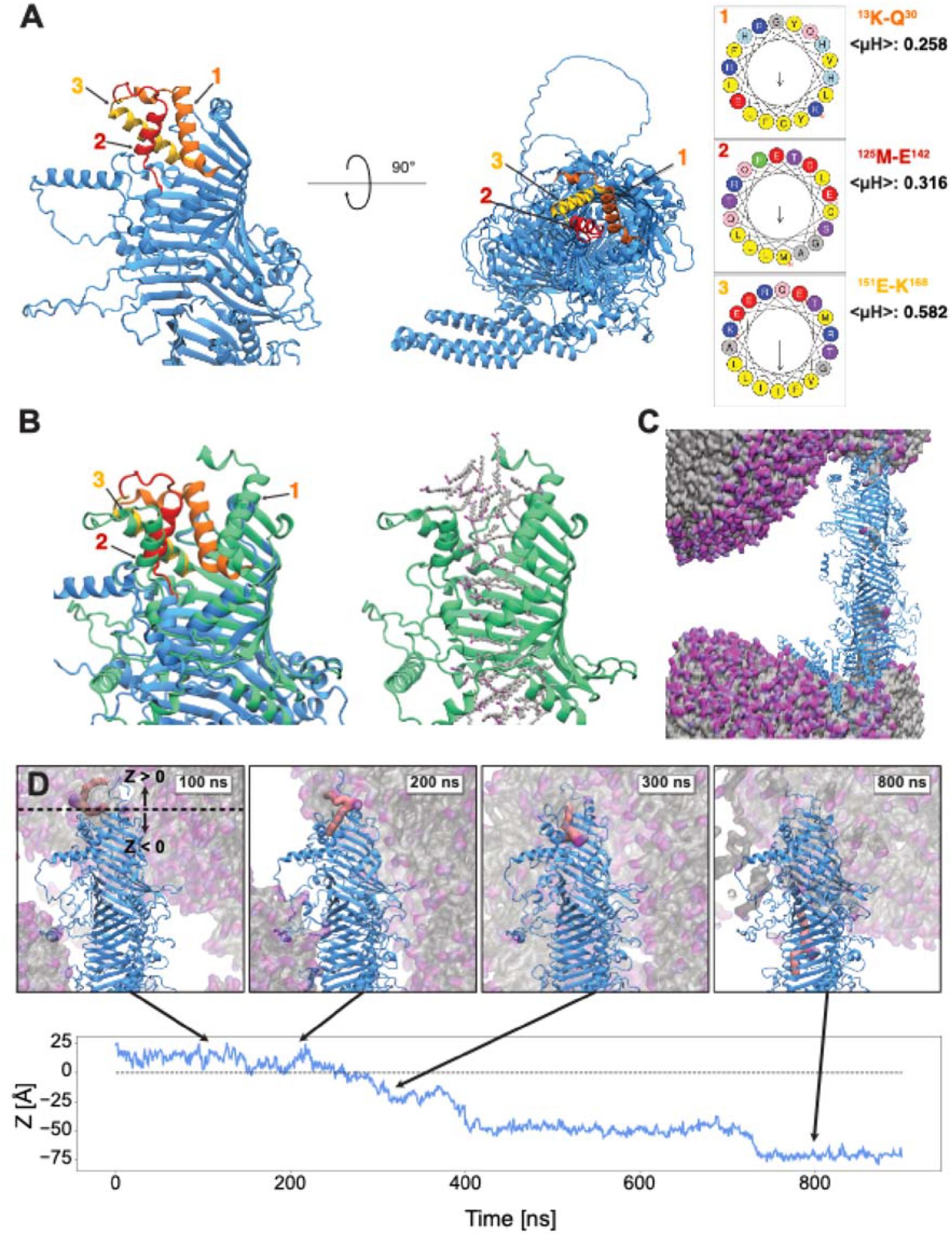
Rearrangement of amphipathic helices enables lipid uptake at ATG2A N-terminus. (A) Amphipathic helices H1-H3 block the hydrophobic cavity of ATG2A at the N-terminus. (right) Heliquest^10^ analysis of helices H1-H3 shows their amphipathic character. (B) AlphaFold 3 prediction of the N-terminal region of ATG2A together with 50 oleic acid molecules shows a displacement of the amphipathic helices (left panel; closed structure as in a, open structure in green), making the cavity accessible to lipids (right panel). (C) The cavity is accessible to lipids (surface representation) from both ATG2A termini (cartoon) in MD simulations (grey: acyl chains, magenta: headgroups). (D) Lipid uptake from the N-terminus. The position of the phosphate head group along the *Z*-axis was tracked, with the entry point of the hydrophobic cavity set at *Z* = 0. All snapshots correspond to CG simulations after backmapping the protein. Disordered loops in the structure are omitted for clarity.

To test this hypothesis, we used AlphaFold 3 to predict the N-terminal fold of ATG2A (residues 1-390) together with 50 molecules of oleic acid. The predicted structure shows a displacement of the amphipathic helices with respect to the initial structure, making the hydrophobic cavity accessible to lipids (Figure 3b, Supplementary Movie 6). H1 shows the biggest displacement, H2 only reorients slightly and H3 barely varies its position. We thus reasoned that H1, as the most flexible due to its position at the N-terminus, may play a role in tethering of ATG2A to the ER membrane.

To test whether the rearrangement of the N-terminus enables lipid uptake and whether the H1 helix contributes to membrane tethering, we performed CG simulations of the full protein with the remodeled N-terminus and dispersed POPC lipids in solution. We found that in these simulations the cavity is accessible to lipids through both termini simultaneously (Figure 3c). Moreover, H1 showed flexibility and bound the head groups of lipids in micelles and vesicles, supporting a potential role of this region in membrane tethering. We observed the same behavior in simulations of ATG2A connecting two parallel bilayers, in which H1 dynamically adjusted its position to interact with the membrane (Supplementary Figure 3).

We noticed that in our model of the closed N-terminus, the positively charged residues in H1 (R15 and H23) directly face negatively charged residues in H2 (E134 and D138, Figure 4a). We thus reasoned that electrostatic interactions between these helices might stabilize the closed conformation. To test this hypothesis, we designed a mutant in which the negative charges in H2 were switched to positive. This should destabilize the H1-H2 interface and hence facilitate the displacement of H1, making the N-terminus accessible and ultimately resulting in a faster lipid transport rate.

**Figure 4.**
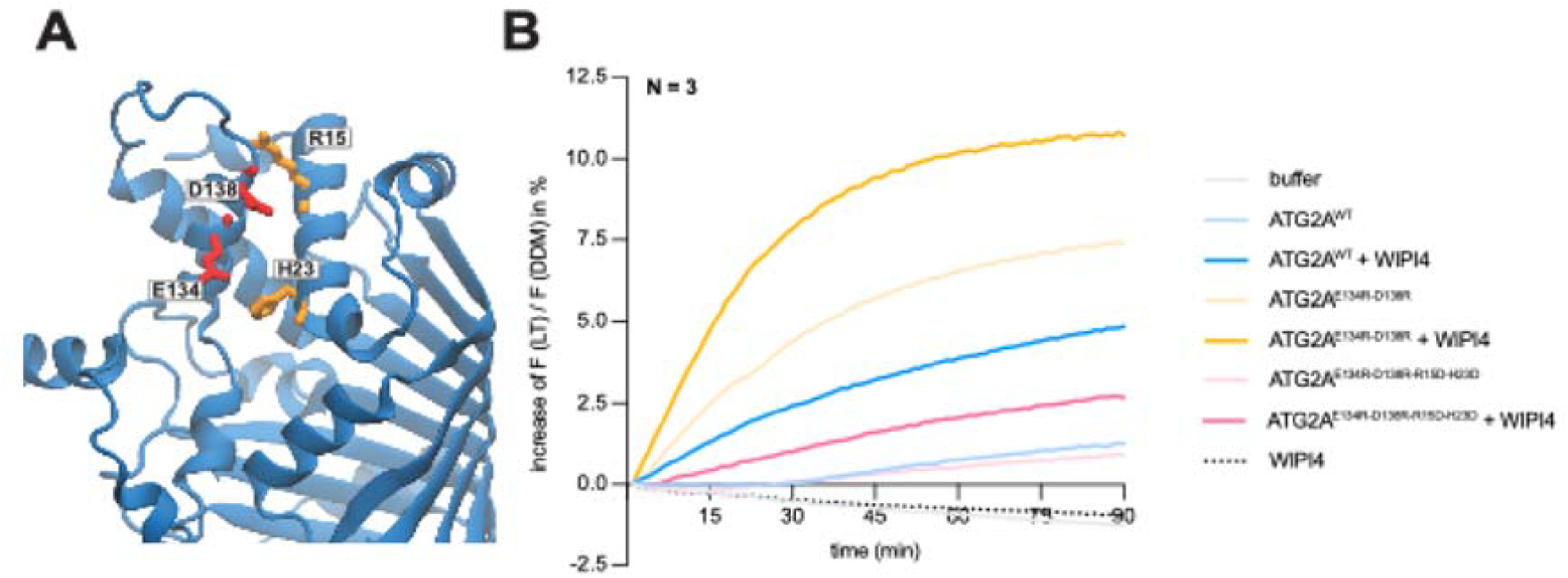
Disruption of the interface between N-terminal helices increases lipid transfer efficiency. (A) Electrostatic interactions stabilize the interface between H1 and H2 helices, blocking the hydrophobic groove (orange: positive residues, red: negative residues). (B) Comparison of lipid transfer activity between wild type, the E134R-D138R mutant, which destabilizes the H1-H2 interface through electrostatic repulsion, and E134R-D138R/R15D-H23D, which recovers electrostatic interactions between H1 and H2. To measure lipid transfer activity, NBD-PE fluorescence was recorded with excitation at 485 nm and emission at 535 nm. The intensity values were normalized by dividing by the maximal fluorescence, recorded 50 minutes after treatment with 0.5% DDM. Addition of WIPI4 increases vesicle tethering and lipid transport efficiency. Lines show the average values of three independent replicas.

We compared lipid transfer activity between the mutant and the WT using a Förster resonance energy transfer (FRET) assay. This method, which is widely used to quantify protein-mediated lipid transport,^20,28,33^ uses donor vesicles containing fluorescent dyes and unlabeled acceptor vesicles. Lipid transport from donor to acceptor vesicles results in dequenching and a subsequent increase in fluorescence intensity. We used 2% of N-(7-nitrobenz-2-oxa1,3-diazol-4-yl)phosphatidylethanolamine (NBD–PE) and L-*α-* phosphatidylethanolamine-N-(lissamine rhodamine B sulfonyl) (Liss Rhod–PE) in the donor liposomes and quantified NBD-PE fluorescence as a measure of lipid transfer activity.

Indeed, although both proteins were used in comparable amounts, as determined by SDS-PAGE gel, the E134R-D138R mutant shows higher lipid transfer efficiency than the wild type (Figure 4b, Supplementary Figure 4). Using a one-rate kinetic model (see Methods), we estimated lipid transport rates of 31.6 lipids/s and 9.4 lipids/s for the D138R-E134R mutant and the wild type, respectively. This represents a 3-fold increase in lipid transfer activity for the mutant, supporting our model of H1 displacement (Supplementary Figure 5). Interestingly, switching the charges of H1 and H2 (D138R-E134R/R15D-H23D), resulted in a decrease in transport efficiency compared to wild type, which is explained by the flipped charge of H1 and the resulting loss of electrostatic interactions with negatively charged membrane. The lower transport efficiency thus supports a role of H1 in membrane tethering (Figure 4b). This agrees with previous results that showed a decrease in transport efficiency for the R15D mutant.^38^ Addition of WIPI4 results in an increase of both vesicle tethering and lipid transport efficiency in the wild type and in both mutants (Figure 4b).

**Figure 5.**
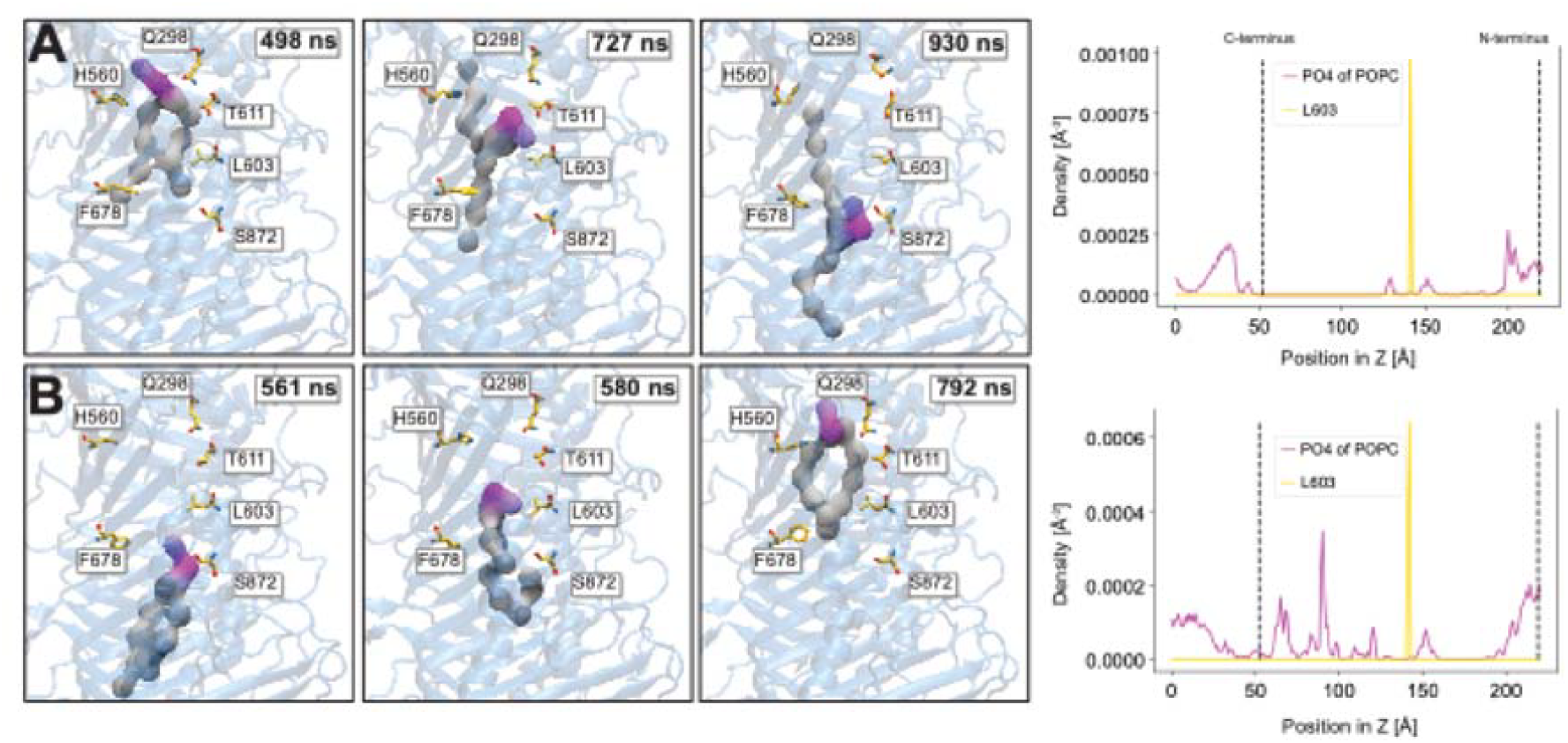
Interactions of lipids at the middle passage region of ATG2A. (A) Snapshots of a simulation where POPC reaches the middle passage region from the N-terminus (left panels) and lipid density at the hydrophobic cavity over simulation time (right panel; *Z* is the position along the principal axis). Polar and charged residues interact with the head groups of the lipid, while hydrophobic residues L603 and F678 interact with the tails. (B) Snapshots and lipid density of a simulation in which POPC arrives to the middle passage from the C-terminus. All snapshots correspond to CG simulations after backmapping the protein.

### 2.4 Lipids are not evenly distributed in the hydrophobic groove of ATG2A

In our MD simulations of ATG2A with displaced N-terminal helices, we observed lipid uptake from vesicles into the hydrophobic cavity through the N-terminus (Figure 3d). Interestingly, although the cavity is accessible from both termini, we never observed a continuously filled channel in our simulations, pointing towards a preference of lipids towards some regions of the cavity. Indeed, the middle passage region only accommodates one lipid in our simulations (Supplementary Figure 6).

**Figure 6.**
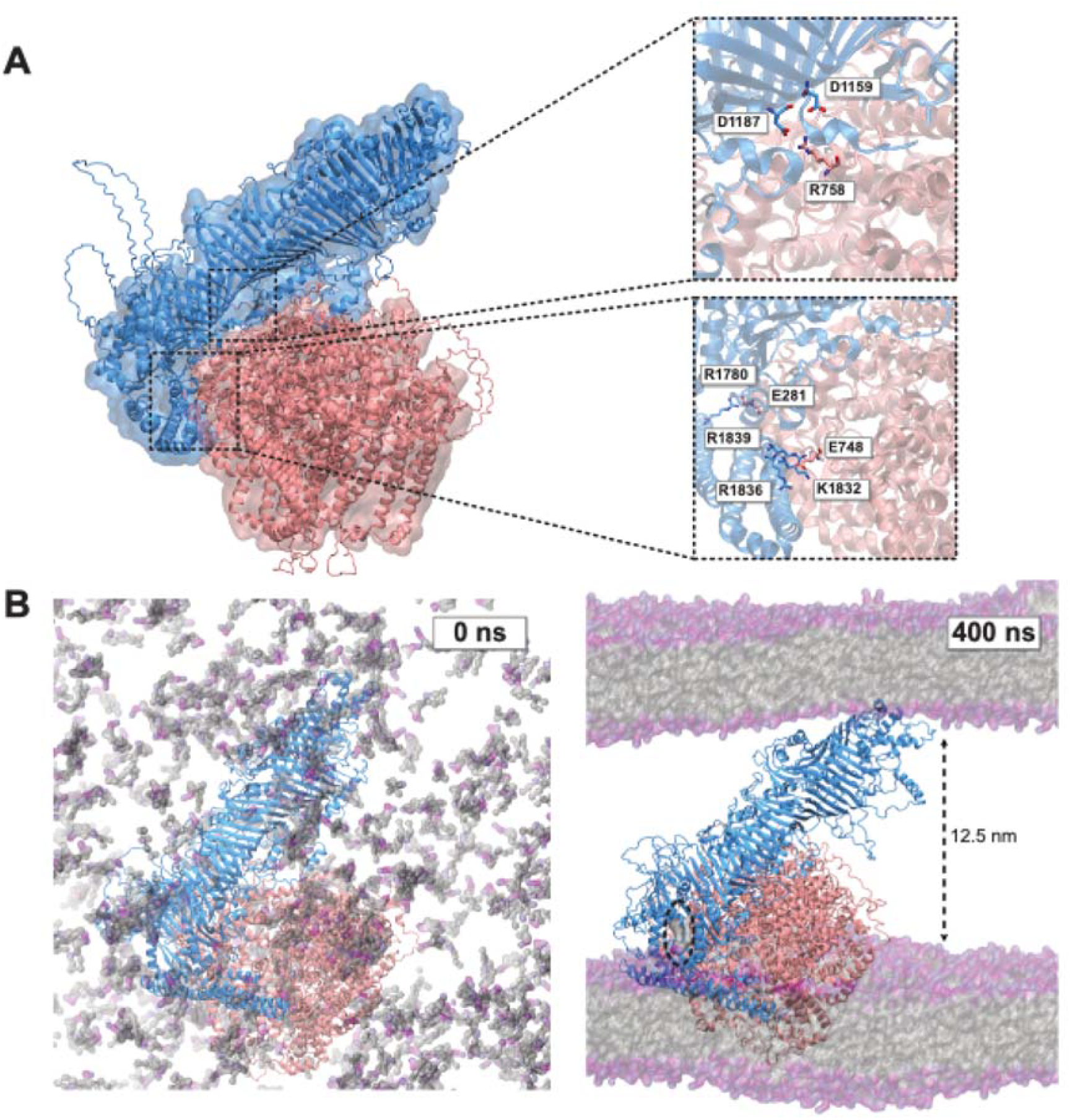
Model of ATG2A-ATG9A complex. (A) AlphaFold 3 prediction of a complex of the ATG9A homotrimer and ATG2A shows an elongated contact interface including charged residues. (B) Spontaneous self-assembly of two parallel lipid bilayers after 400 ns of CG simulation of the ATG2A-ATG9A complex with initially dispersed POPC lipids. The hydrophobic cavity of ATG2A is accessible from the membrane at the C-terminus and contains one lipid (dashed ellipse). See Supplementary Movie 9 for the spontaneous formation of the two membranes connected by ATG2A anchored to ATG9A.

In two replicas, we saw “jumps” of lipids from the N- and C-terminus to the middle passage region between residues L603 and F678, where the lipids stay stably bound for hundreds of nanoseconds (Supplementary Movies 7 and 8). This observation is consistent with our simulation of the N-terminally inaccessible ATG2A described above, where a lipid was retained in the middle passage region. Wang et al.^38^ suggest that lipid reorientation might be required for passage through this region. Interestingly, in the simulation in which the lipid entered through the N-terminus and got trapped between F678 and L603 on its way down, the lipid underwent a reorientation in which one tail and the head group passed through the gate (Supplementary Movie 7, Figure 5).

Upon reorientation of the lipid in the middle passage region, there is no steric hindrance for it to move towards the C-terminus. We therefore attribute lipid trapping to strong interactions with a cluster of residues at this site. Indeed, a closer look shows that while the head group interacts with polar and charged residues such as Q298, H560, T611, and S872, the hydrophobic tails stably engage with L603 and F678 (Figure 5).

### 2.5 Model of ATG2A-ATG9A interaction

Over the past years, several efforts to elucidate the structure of the ATG2A-ATG9A complex have been reported. So far, two different models have been proposed, both of which involve a direct lipid transfer from ATG2A to ATG9A, but differ in the delivery sites (i.e., the perpendicular branch vs. the lateral pore).^34,38^ Our simulations indicate that ATG2A can deliver lipids directly to micelles through the C-terminus. This agrees with *in vitro* data, which show that Atg2 is sufficient to transport lipids to liposomes in the absence of Atg9.^20,28,33^

Interestingly, AlphaFold 3 predicts a complex between ATG2A and ATG9A in which the interaction interface of ATG2A does not include the opening of the hydrophobic cavity, but rather the back side of the hydrophobic groove. The predicted interaction interface is weak but elongated, including residues from the C-terminal region and the more N-terminal part of the protein (Figure 6a).

We thus wondered whether the AlphaFold 3 model is compatible with the position and orientation of both proteins in the lipid bilayer. To answer this question in an unbiased manner, we ran MD simulations of the complex with POPC lipids initially dispersed in solution. Interestingly, in one replicate two parallel membranes assembled, one of them encapsulating the well-characterized transmembrane region of ATG9A and interacting with the C-terminus of ATG2A, and the other interacting with its N-terminus (Supplementary Figure 7b; Supplementary Movie 9). Other replicates showed formation of vesicles containing ATG9A and tethered by ATG2A (Supplementary Figure 7a), again consistent with lipid transfer function.

In the predicted complex, the N-terminus of ATG2A is not accessible to lipids, with the aforementioned helices blocking the entry of the hydrophobic groove. We thus modeled this region separately and repeated the lipid self-assembly simulations to elucidate whether this region has an impact on membrane orientation. In these simulations, lipids assembled in a similar manner to form two parallel bilayers, suggesting that the open conformation is also compatible with this orientation of the complex (Supplementary Figure 7b).

To assess the compatibility of this putative ATG2A-ATG9A complex with WIPI4 binding, we superimposed the groove of ATG2A in our model onto that of the cryoEM structure of the ATG2A-WIPI4 complex resolved by Wang et al..^38^ We aligned the structure in the membrane-bound complex (Figure 5b) to elucidate whether the ATG2A-ATG9A-WIPI4 complex would be compatible with this membrane orientation. The region interacting with WIPI4, including residues L1578, R1579 R1668, T1670, R1711, and L1709, is unoccupied in our complex and there were no clashes with ATG9A upon superimposition. Moreover, the LRRG motif of WIPI4, which is known to interact with PI3P, is close enough to the membrane that slight reorientations would allow its interaction (Supplementary Figure 8). This suggests that our ATG2A-ATG9A model is not only compatible with the position and orientation of ATG2A and ATG9A in the membrane, but also with the interaction of the complex with WIPI4.

## 3 Discussion

We have combined structural predictions, MD simulations, and lipid transfer assays to shed light onto the mechanism of lipid transport by ATG2A. Our results indicate that ATG2A can bind to membranes through both termini simultaneously and support bidirectional lipid transport. We propose that a rearrangement of amphipathic helices at the N-terminus is necessary for lipid uptake. Supporting this model, mutations facilitating this reorientation strongly enhance lipid transport *in vitro*. Finally, our data point towards an uneven distribution of lipids within the cavity, with a narrow N-terminus, unoccupied sites in the middle passage region and a reservoir of lipids at the C-terminus.

Our findings also support previous experimental work suggesting that ATG2A can tether membranes by binding to lipids through both termini simultaneously. We were able to reproduce the membrane orientation and intermembrane distances observed using negative-stain EM and *in situ* cryo-electron tomography. Moreover, our simulations are consistent with *in vitro* data indicating a preferential binding of ATG2A to highly curved membranes. Notably, we observed lipid transport events in the microsecond time range in unbiased simulations. This would be compatible with the transport rates that have been estimated from lipid transfer fluorescence assays of 17100 lipids per second.^20,36^

Insights from our work may clarify unresolved aspects of Atg2-mediated lipid transport. The necessary rearrangements of the amphipathic N-terminal helices may account for the high flexibility in this region, which remained unresolved in the cryo-EM structure reported by Wang et al..^38^ These reorientations might also explain the slow phase of lipid transport rates from fluorescence assays, reflecting a small Atg2 subpopulation in which the helices were not fully displaced.^36^ Interestingly, Wang et al.^38^ have shown that mutations R15D and K13D at the H1 helix reduce lipid transport, further supporting a putative role of this helix in membrane tethering. Finally, the uneven distribution of lipids in the cavity during our simulations might point towards a selectivity filter, although more work in this direction would be required to support this claim.

Two different lipid transport mechanisms for Atg2 have been proposed: the *bridge* and the *ferry* model.^26^ In the bridge model, Atg2 transports lipids between tethered membranes, while in the ferry model, Atg2 loads lipids from one membrane, unbinds, and delivers them to another membrane. Our simulations and *in vitro* transport assays suggest that both mechanisms are possible, but the bridge is more efficient. However, overexpression of an N-terminal fragment of Atg2, which is not able to bridge two membranes completely, recovered autophagy in ATG2 knockout cells.^33^ This indicates that the ferry mechanism is, at least under overexpression, sufficient for autophagosome formation. Our findings suggest that the N-terminal H1 helix might act as a plug and mediate opening and closing of the hydrophobic cavity. We hypothesize that such a gating mechanism only makes sense upon simultaneous binding of both membranes, hence supporting the bridge transport model. Future work in this direction could include simulations of membrane assembly with the N-terminal Atg2 fragment.

We propose an ATG2A-ATG9A complex that, unlike other recent models,^34,38^ does not involve direct lipid handover from ATG2A to ATG9A. In our model, the hydrophobic cavity of ATG2A is in direct contact with the membrane and can thus deliver lipids directly to the membrane. This might seem contradictory to experiments reported by Gomez-Sanchez et al.,^12^ who identified point mutations at the entry of the hydrophobic cavity that impair Atg2-Atg9 interactions in yeast. Although these observations make the model of direct lipid handover very appealing, such a model would imply that the delivery process depends on Atg9. In vitro fluorescence transport assays show, however, that Atg2 is sufficient for efficient lipid transport. While the mutations proposed by Gomez-Sanchez et al.^12^ are not at the ATG2A-ATG9A interface in our model, it cannot be excluded that their effects on the interaction are indirect. Based on our simulations, we propose that the interaction with ATG9A might be important for correct localization and orientation of Atg2 as well as the subsequent equilibration of the transported lipids between the monolayers, without being strictly required for lipid transport *per se*.

All in all, our findings provide valuable insights into Atg2-mediated lipid transport. Due to the variety of cellular processes in which Atg2 is involved, such as lysosomal repair, understanding its mechanistic details is not only relevant in the context of autophagy. Furthermore, our findings might shed light on the mechanisms of other lipid transporters, such as VPS13. More broadly, Atg2 could serve as a model for studying organelle contact sites.

## 4 Methods

### 4.1 Molecular dynamics (MD) simulations

#### 4.1.1 System preparation

We used AlphaFold 3 to generate structures of ATG2A and the ATG2A-ATG9A complex.^2^ To obtain a model of the open N-terminus of ATG2A, we included 50 molecules of oleic acid in the prediction of the N-terminal region (residues 1-390). *Martinize2* was used to map the atomistic models to CG resolution.^17^ To maintain the secondary structure of the ordered regions of the protein, we used an elastic network with a force constant of 500 kJ mol^−1^ nm^−2^ and lower and upper cutoffs of 0.5 and 0.9 nm, respectively. POPC lipids were placed at random positions at different concentrations (Supplementary Table 1). The systems were solvated with water and 150 mM of NaCl.

#### 4.1.2 Unbiased membrane binding from CG simulations of lipid self-assembly

All simulations were run using Martini 3 and Gromacs 2022.4.^1,31^ In some systems, the C1 beads of POPC were modified to C1r to reduce self-interactions (Supplementary Table 1). We minimized the potential energy of the system using the steepest descent algorithm for 50,000 steps. We then performed the lipid self-assembly simulations in the NPT ensemble using the Verlet cutoff scheme with cut-off distance of 1.418 nm for the short-range neighbor list, which was updated every 20 steps. Coulomb and van-der Waals interactions were cut-off at 1.2 nm. A relative dielectric constant of 15 was used. Temperature was kept constant at 310 K using the Berendsen thermostat^4^ and a coupling constant of 2 ps. Pressure was kept constant at 1 bar isotropically using the Berendsen barostat and a coupling constant of 6 ps, with compressibility of 3 × 10^−4^ bar^−1^. A detailed protocol is available here: dx.doi.org/10.17504/protocols.io.dm6gpm1j1gzp/v1.

#### 4.1.3 Simulations of the C-terminus of ATG2A in a buckled bilayer

To investigate the curvature preference of ATG2A, we first generated a POPC bilayer using the insane^39^ method. To buckle the bilayer, we ran a short 10 ns simulation using the Berendsen barostat, coupled anisotropically to the *Y* and *Z* directions, with a nonzero compressibility of 3 × 10^−4^ bar^−1^ only in these dimensions. The applied pressure was set to 12 bar in the *Y* direction and 1 bar in Z. We then ran a simulation of the buckled membrane with the C-terminus of ATG2A initially placed at a random position in the simulation box. To keep the membrane curvature constant, we fixed the lateral axes by coupling the Parrinello-Rahman barostat^29^ only to the *Z* dimension, with a compressibility of 3 × 10^−4^ bar^−1^ and a target pressure of 1 bar. After 1 *µ*s of simulation the C-terminus had stably bound the bilayer. We used this as a starting configuration for 10 independent replicas with different initial velocities drawn independently from the Maxwell-Boltzmann distribution. A detailed protocol is available here: dx.doi.org/10.17504/protocols.io.3byl468jogo5/v1.

#### 4.1.4 Calculation of local membrane curvature

Local membrane curvature was calculated using a 2D Fourier expansion as has been described previously,^5^ using the publicly available code (https://github.com/bio-phys/MemCurv/tree/master). To calculate the lowest *k* = 3 Fourier coefficients, we used the position of each POPC phosphate group (PO4 beads) at 1 ns intervals along the trajectory. We used the computed height profile to calculate the local curvature of the membrane at the position of the center of mass of the C-terminus of Atg2. For reference, we performed the same calculation for randomly selected lipids, which were newly chosen every saved frame.

#### 4.1.5 Lipid uptake and release

To track individual lipids and describe their position in the hydrophobic cavity of ATG2A, we first fixed the protein and aligned the simulation box to it. To describe entry and release events through the C-terminus, residue S1791 was used as a reference. Hence, the *Z* component of its position was subtracted from the *Z* component of the position of the given lipid, making positive values correspond to the interior of the groove. To describe lipid uptake through the N-terminus, residue E151 was used as reference.

#### 4.1.6 Lipid density calculation

We used the DensityAnalysis class from the MDanalysis library to calculate lipid densities at the hydrophobic cavity of Atg2.^13,22^ To restrict the density calculation to the cavity, we used a grid of 30 Å × 30 Å × 220 Å at 1 Å mesh width placed around the center of mass of the protein.

#### 4.1.7 Visualization

We used VMD 1.9.4 for visualization and the Tachyon ray-tracer for image generation.^14,32^ For visualization purposes, we used the CG2AT software to backmap the protein into atomistic resolution in all simulation snapshots included in this manuscript.^35^

### 4.2 Purification of ATG2A and Mutant Variants from HEK293F Cells

Recombinant 3XFLAG-TEV-tagged ATG2A and its mutants were expressed from a pcDNA3.1(+) vector (RRID:Addgene 244949). Mutations were introduced via site-directed mutagenesis. Constructs were transiently transfected into FreeStyle^*™*^ HEK293F cells (Thermo Fisher Scientific) maintained at 37 °C in FreeStyle^*™*^ 293 Expression Medium (Cat# 12338-026). Cells were seeded at a density of 0.7 × 10^6^ cells/mL one day before transfection. On the day of transfection, 400 *µ*g of plasmid DNA (MAXI prep) was mixed with 13 mL Opti-MEM^*™*^ I Reduced Serum Medium (Cat# 31985-062) and combined with 800 *µ*g PEI 25K (Polysciences, Cat# 23966-1), also diluted in 13 mL of Opti-MEM. This mixture was added to a 400 mL culture. Twenty-four hours post-transfection, 100 mL of EX-CELL^®^ 293 Serum-Free Medium (Sigma-Aldrich, Cat# 14571C) was added to support protein production. After 72 hours, cells were collected by centrifugation at 270 × g for 20 minutes, washed once with PBS, and flash-frozen in liquid nitrogen for storage. For protein purification, frozen pellets were thawed on ice in 20 mL of lysis buffer (50 mM HEPES pH 8.0, 300 mM NaCl, 10% glycerol, 1 mM TCEP, 0.5 M urea, 0.2% n-octyl*β*-D-glucopyranoside (OG; Glycon, Cat# D97001-10g)), supplemented with cOmplete^*™*^ EDTA-free protease inhibitors (Roche), Benzonase (1 *µ*L/50 mL, Sigma-Aldrich, Cat# E1014), and Protease Inhibitor Cocktail (200 *µ*L/50 mL, Sigma-Aldrich, Cat# P8849). Cells were lysed using a Dounce homogenizer (10 strokes with the loose pestle followed by 2 × 20 strokes with the tight pestle), and lysates were clarified by centrifugation at 15,000 rpm for 30 minutes at 4 ?C (HITACHI centrifuge). Clarified lysates were incubated with 0.6 mL of pre-equilibrated anti-FLAG M2 resin (Sigma, Cat# A2220) for 3 hours at 4 °C with gentle rotation. The resin was washed three times with wash buffer containing glycerol (50 mM HEPES pH 8.0, 300 mM NaCl, 10% glycerol, 1 mM TCEP) and an additional three times with glycerol-free buffer of the same composition. Beads were transferred to microcentrifuge tubes and incubated overnight at 4 °C with 200 µL of glycerol-free wash buffer supplemented with 5 *µ* L TEV protease for on-bead cleavage. The next morning, the eluate (Elution 1) was collected, and residual protein was recovered by washing beads with 150 *µ*L of wash buffer without glycerol (Elution 2). All purification steps were performed at 4 °C or on ice. OG detergent (CMC ≈ 0.6%) was added from a 10% aqueous stock to aid solubilization of membrane-associated protein species. A detailed protocol is available (dx.doi.org/10.17504/protocols.io.n92ld6zdng5b/v1).

### 4.3 Protein expression and purification of WIPI4 from insect cells

A codon-optimized synthetic gene for WIPI4 (Addgene 244943) was cloned into a baculoviral expression vector and expressed in Sf9 insect cells. Cells were harvested, lysed, and the protein was purified via a HisTrap HP column (Cytiva, Cat# 17524802) followed by size-exclusion chromatography (Superdex 200 Increase 10/300 GL column, Cat# 28990944). Purified protein was aliquoted, snap-frozen in liquid nitrogen, and stored at -80 °C. A detailed step-by-step protocol is available at protocols.io (dx.doi.org/10.17504/protocols.io.81wgbw6bqgpk/v1).

### 4.4 Liposome preparation

Small unilamellar vesicles (SUVs) were prepared using the lipid compositions described below. Lipids dissolved in chloroform were mixed in glass vials, evaporated under a gentle argon stream, and dried further under vacuum for 1 hour to remove residual solvent. The dried lipid films were rehydrated in SEC300 buffer (25 mM HEPES pH 7.5, 150 mM NaCl, 1 mM TCEP), gently vortexed, and sonicated in a bath sonicator for 2 minutes. The suspension was subsequently extruded through a 0.1 *µ*m membrane filter (Whatman, Cat# 10419504) using an Avanti Mini Extruder (Cat# 610024). The final SUV suspension had a lipid concentration of 0.5 mg/mL. A detailed protocol is available (dx.doi.org/10.17504/protocols.io.x54v95y6ml3e/v1).

Donor liposomes were composed of 66% DOPC, 30% DOPE, 2% NBD-PE, and 2% Rh-PE. Acceptor liposomes contained 65% DOPC, 15% DOPE, 15% DOPS, and 5% PI3P. The lipids used in this study included: DOPC (Avanti, Cat# 850375C), DOPE (Avanti, Cat# 850725C), DOPS (Avanti, Cat# 840035C), PI3P (Sigma-Aldrich, Cat# 850150P500UG), NBD-PE (Sigma-Aldrich, Cat# 810155P-1MG), and Rh-PE (Invitrogen, Cat# L-1392).

### 4.5 Lipid transfer assay

Lipid transport was monitored using 384-well glass-bottom plates (Greiner Bio-One, Cat# 781892) at room temperature. Each reaction (50 *µ*L total volume) contained 2.5 *µ*L each of donor and acceptor liposomes (0.5 mg/mL lipid), along with recombinant protein as indicated in figure legends (250 nM ATG2A), all in SEC300 buffer. NBD fluorescence was recorded every 20 seconds for 90 minutes using a Tecan SPARK plate reader (RRID:SCR 021897; Tecan Life Sciences), with excitation at 485 nm and emission at 535 nm. To determine the maximal fluorescence, 0.5% (v/v) DDM was added to solubilize all membranes, and fluorescence was measured 50 minutes post-DDM addition. Fluorescence data were normalized by subtracting the baseline signal and dividing by the DDM-treated endpoint value.

To assess whether ATG2A variants contain similar protein concentrations, samples were loaded onto 4–12% SDS-PAGE gels (Invitrogen, Cat# NP0321BOX or NP0323BOX) with PageRuler Prestained Protein Marker (Thermo Fisher, Cat# 26620 or 26617).

### 4.6 Estimation of lipid transfer rates from experimental data

To fit the *in vitro* lipid transfer data (observed fluorescence), we used the theory previously developed by von Bülow and Hummer.^36^ The observed fluorescence is described as a one-rate kinetic model of the following form,

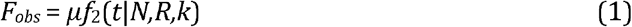

given a number of lipids *N* and vesicle size *R*, with *k* and *µ* denoting rate of transfer and fluorescence amplitude, respectively. Here, *f*_*2*_ is defined as a scaled version of the expected fluorescence intensity *F*_*2*_:

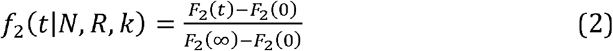

Where

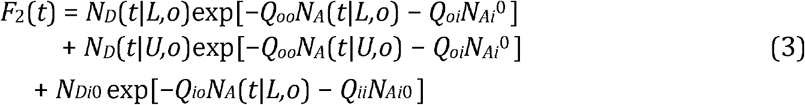

Here, *D* and *A* denote donor and acceptor lipids, *L* and *U* labeled and unlabeled vesicles, and *i* and *o* inner and outer leaflets. Intra- and interleaflet quenching factors *Q*_*ij*_ were used as described in Ref. 36. Note that we neglect the effect of lipid flip-flop in this model. The populations *N*_*i*_ as function of time *t* were modeled as in Ref. 36. Since we used a 100 nm filter for liposome extrusion, we assumed all vesicles to have a radius of *R* = 50 nm.

## 5 Acknowledgements

We thank Martina Schuschnig for help with cloning the ATG2A constructs, Sanjoy Paul and Dorotea Fracchiolla for insightful discussions, and the Vienna BioCenter Core Facilities (VBCF) Protech Facility for help with HEK cell expressions.

## 5.3 Data availability

The data, code, protocols and key lab materials used and generated in this study are listed in a Key Resource Table alongside their persistent identifiers at https://doi.org/10.5281/zenodo.17507215. This repository also includes parameter files and starting structures used for simulations, compressed trajectories, and high-resolution versions of all supplementary movies.

