## Supplementary Information for "Pathway and gates for ATG2A-mediated lipid transport in autophagy"

**Supplementary Table 1**: Summary of all simulations used in this study

| Protein | Lipid environment | Box size ($A^3$) | C1 beads in POPC* | Position restraints | Simulation length | # Replicates | Total simulation time |
|---|---|---|---|---|---|---|---|
| **ATG2A$^{full\text{-}length}$ N-ter closed** | 250 POPC in solution | 15 * 15 * 21 | C1r | Yes** | 400 ns | 10 | 4000 ns |
| **ATG2A$^{full\text{-}length}$ N-ter closed** | 500 POPC in solution | 15 * 15 * 21 | C1r | No | 410-3197 ns | 10 | 24446 ns |
| **ATG2A $^{C\text{-}ter}$ (residues 1-239)** | Buckled POPC membrane | 20.7 * 18.8 * 19.5 | C1 | No | 1000 ns | 10 | 10000 ns |
| **ATG2A$^{full\text{-}length}$ N-ter open** | 2500 POPC in solution | 25 * 25 * 25 | C1 | Yes* | 1000 ns | 20 | 20000 ns |
| **ATG2A-ATG9A N-ter closed** | 2500 POPC in solution | 25 * 25 * 25 | C1 | Yes* | 400 ns | 10 | 4000 ns |
| **ATG2A-ATG9A N-ter open** | 2500 POPC in solution | 25 * 25 * 25 | C1 | Yes* | 1000 ns | 10 | 10000 ns |

* The C1 beads in the POPC tails were either kept as in the original force field (C1) or changed to C1r, resulting in a reduction of the self-interactions

** Except in flexible loops and c-terminus of ATG2A (A1700-D1938)

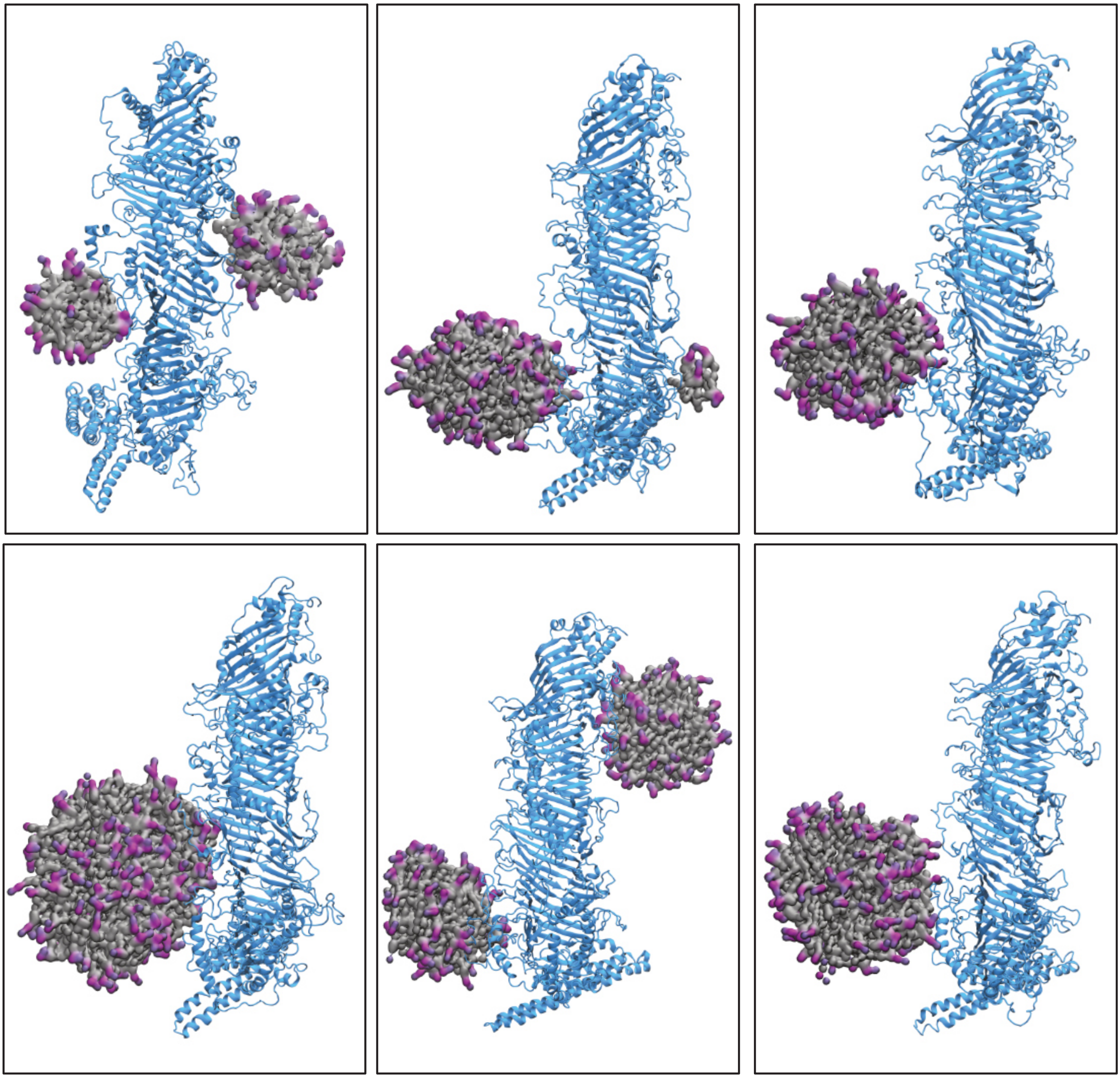

**Supplementary Figure 1: Lipid aggregates transiently interact with disordered regions at the surface of ATG2A.** All snapshots correspond to CG simulations of different replicates after backmapping the protein.

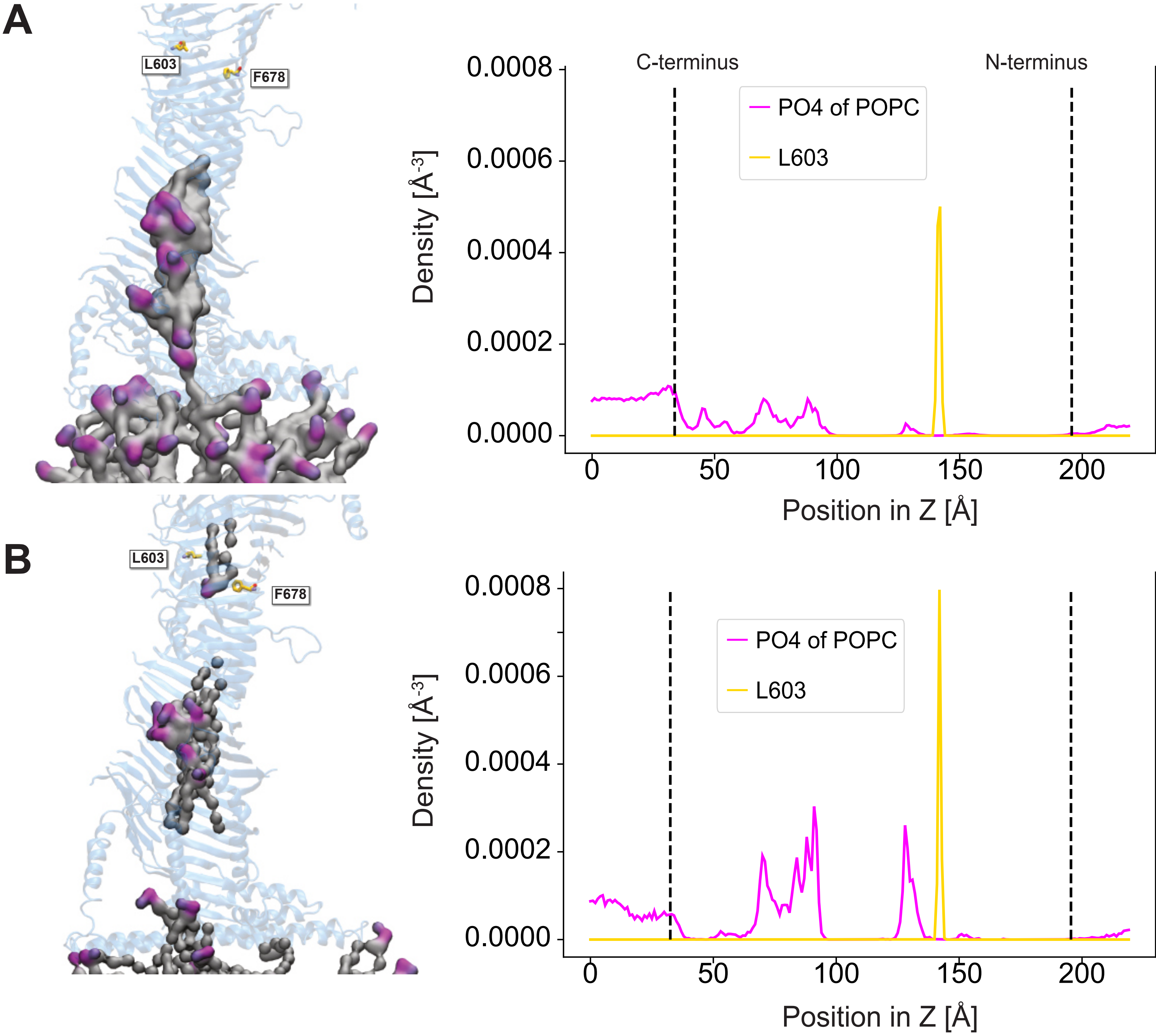


**Supplementary Figure 2: The N-terminal region of the hydrophobic cavity is not accessible for lipids entering through the C-terminus.** (A) Representative snapshot of lipids occupying only the C-terminal ~6.5 nm of the hydrophobic groove (left) and average density of lipids in the cavity for 10 replicates (right). Lipid density was calculated along the direction Z of the long axis of ATG2A defined by its inertia tensor. To restrict the density calculation to the cavity, we used a grid of 30 Å x 30 Å x 220 Å at 1 Å mesh width placed around the center of mass of the protein. (B) Snapshot and lipid density of a simulation in which one lipid was "trapped" at the middle passage region. All snapshots correspond to CG simulations of different replicates after backmapping the protein. Disordered loops in the structure are omitted for clarity.

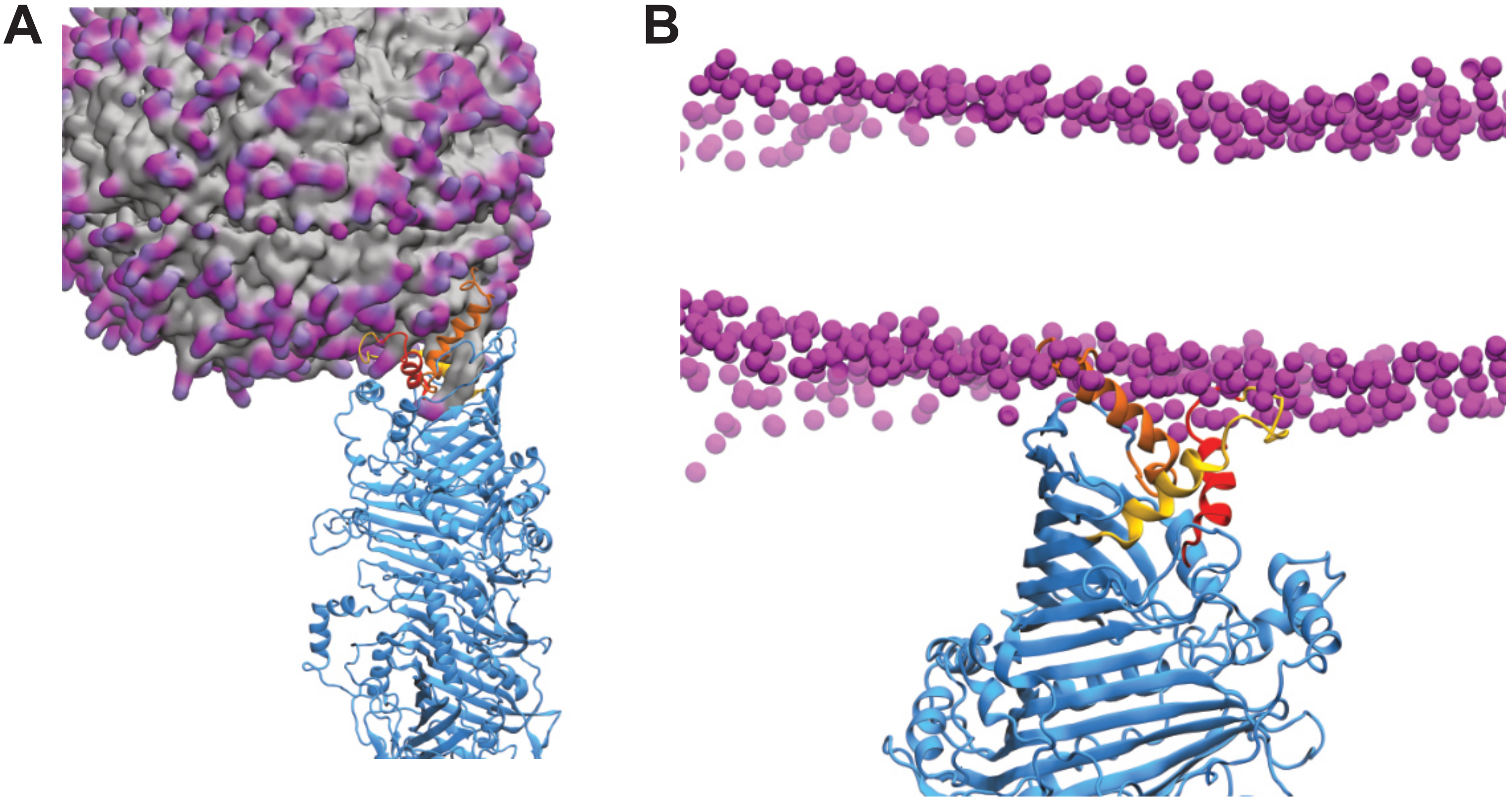


**Supplementary Figure 3: Amphipathic N-terminal helices tether ATG2A to lipid membranes.** Snapshots of N-terminal tethering to a POPC vesicle (A) and a POPC bilayer (B). Both snapshots correspond to CG simulations after backmapping the protein.

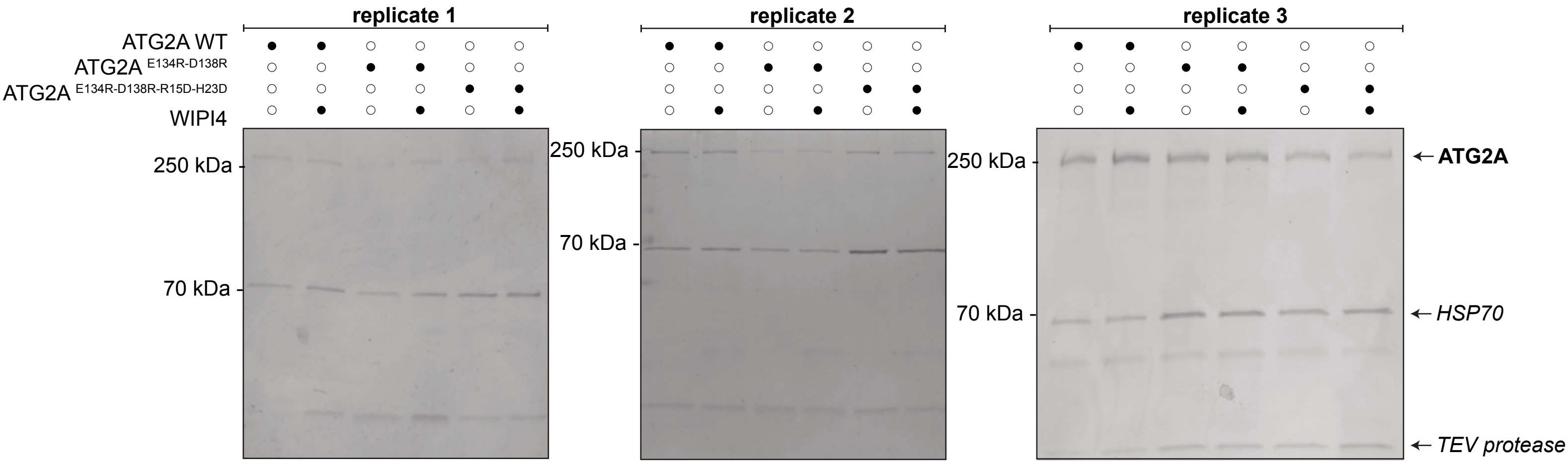


**Supplementary Figure 4: SDS-Page gel shows that protein levels used for the lipid transfer FRET assay are comparable.**

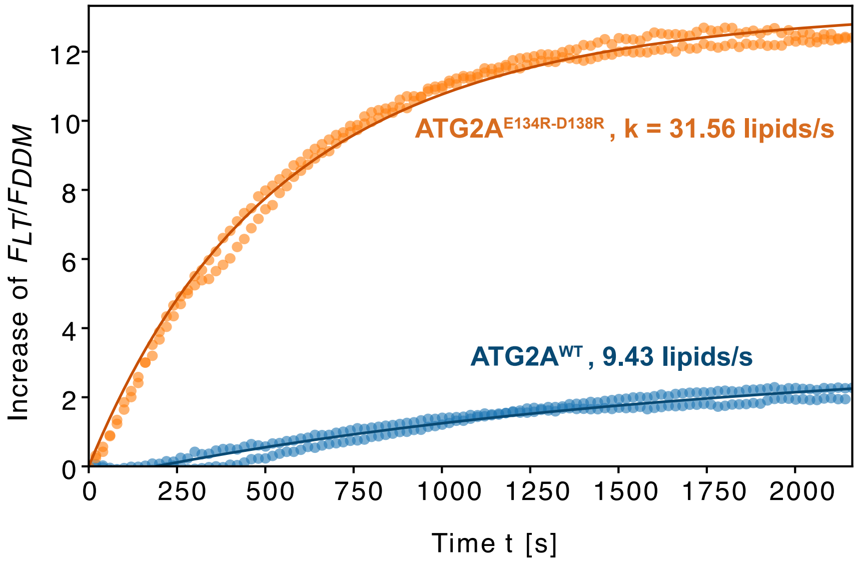


**Supplementary Figure 5: Disruption of the H1-H2 interface through electrostatic repulsion leads to a 3-fold increase in lipid transfer efficiency.** Data points from two independent replicates were fitted together (solid lines) using a one-rate kinetic model (see Methods). To fit the kinetic model of the WT protein, an offset of 200 seconds was used.

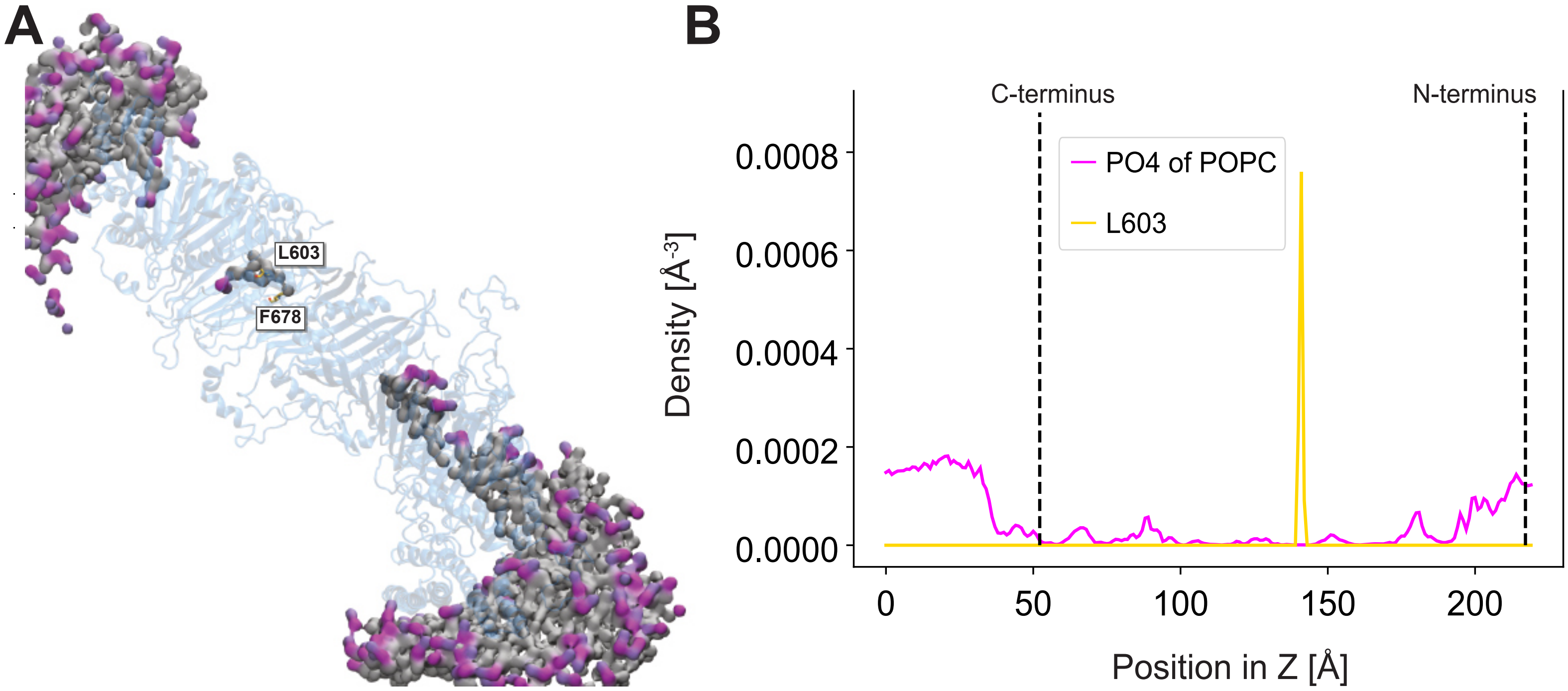


**Supplementary Figure 6: Lipids do not form a continuous channel at the hydrophobic cavity of ATG2A, even when both termini are accessible from solution.** (A) Representative snapshot of lipids occupying only the C-terminal ~6.5 nm of the hydrophobic groove, the middle passage region and the N-terminal end of the cavity. The snapshot corresponds to a CG simulation after backmapping the protein. (B) Average density of lipids in the cavity for 20 replicates, each 1 μs long. Lipid density was calculated along the direction Z of the long axis of ATG2A defined by its inertia tensor. To restrict the density calculation to the cavity, we used a grid of 30 Å x 30 Å x 220 Å at 1 Å mesh width placed around the center of mass of the protein.

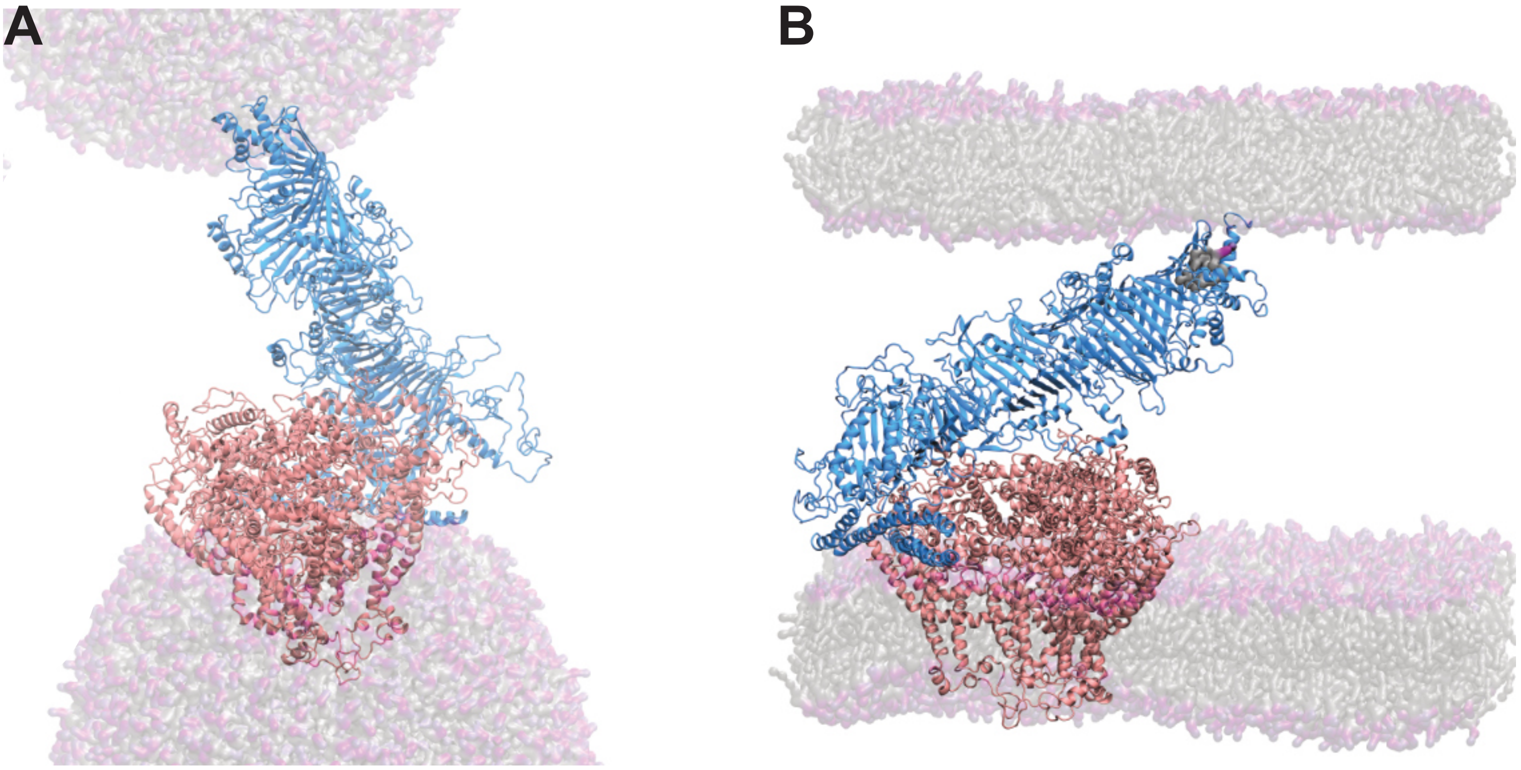


**Supplementary Figure 7: Snapshots of the ATG2A-ATG9A complex tethering two vesicles (A) and two parallel lipid bilayers (B).** The snapshots correspond to CG simulations after backmapping the proteins.

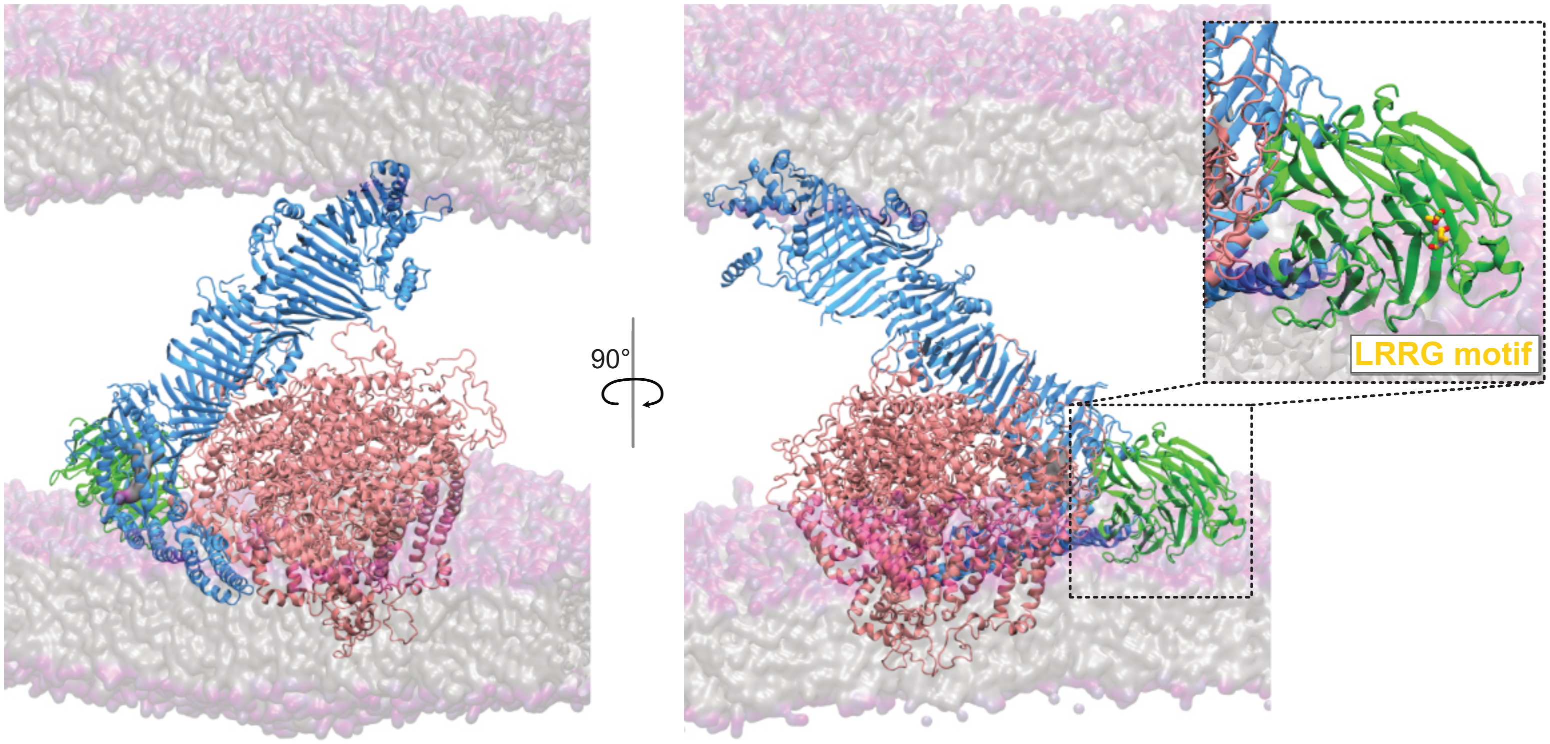


**Supplementary Figure 8: ATG2A-ATG9A complex is compatible with WIPI4 binding.** Snapshot of a simulation after superimposition of the groove of ATG2A in our model with the one in the cryo-EM structure of the ATG2A-WIPI4 complex resolved by Wang et al. (PDB: **8KBX**). WIPI4 does not clash with ATG9A and the LRRG motif is just above the membrane and accessible to binding by phosphoinositol PI(3)P lipids. Disordered loops in the structure of ATG2A are omitted for clarity.
